# Wild Steps in a Semi-wild Setting? Habitat Selection and Behavior of European Bison reintroduced to an Anthropogenic Landscape

**DOI:** 10.1101/325985

**Authors:** Pil Birkefeldt Møller Pedersen, Joanna B. Olsen, Brody Sandel, Jens-Christian Svenning

**Affiliations:** Section for Ecoinformatics & Biodiversity, Department of Bioscience, Aarhus University, Aarhus, Denmark; Center for Biodiversity Dynamics in a Changing World (BIOCHANGE), Aarhus University, Aarhus, Denmark

## Abstract

Recently, several wild or semi-wild herds of European bison have been reintroduced across Europe. It is essential for future successful bison reintroductions to know how the European bison use different habitats, which environmental parameters drive their habitat selection, and whether their habitat use and behavioural patterns in new reintroduction sites differ from habitats where European bison have been roaming freely for a long time.

Here, we address these questions for a 40-ha enclosed site that has been inhabited by semi-free ranging European bison since 2012. The site, Vorup Meadows, is adjacent to the Gudenå river in Denmark and consists of human-modified riparian meadows. During 2013 we monitored the behavioural pattern and spatial use of the 11 bison present and in parallel carried out floristic analyses to assess habitat structure and food quality in the enclosure. We tested habitat use and selection against environmental parameters such as habitat characteristics, plant community traits, topography, and management area (release area vs. meadow area) using linear regression and spatial models.

The bison herd had comparable diurnal activity patterns as observed in previous studies on free-roaming bison herds. Topography emerged as the main predictor of the frequency of occurrence in our spatial models, with high-lying drier areas being used more. Bison did not prefer open areas over areas with tree cover when accounting for habitat availability. However, they spent significantly more time in the release area, a former agricultural field with supplementary fodder, than expected from availability compared to the rest of the enclosure, a meadow with tree patches. We wish to increase awareness of possible long-term ethological effects of the release site and the management protocols accomplished here that might reduce the ecological impact by the bison in the target habitat, and thereby compromise or even oppose the conservation goals of the conservation efforts.

## 1. Introduction

In the last few decades, European bison (*Bison bonasus*) have been reintroduced to various habitats across Europe, ranging from coastal dunes in the Netherlands (e.g. 1, 2) over mountainous mosaic landscapes in the French Alps (3)and in Germany (4) to lowland peri-urban meadows (5). Reintroduction areas not only represent different habitats, they also represent different degrees of human modification, with some reintroduction sites being naturally disturbed coastal dunes (e.g. 2), private commercial plantation (e.g. 4) or areas still ditched and drained due to upland agricultural use (5). The conservation goal of these reintroductions is often twofold, with one focus being the protection of the largest extant wild, native herbivore in Europe, European bison (6), and the other focus being ecological restoration focusing on restoring trophic top-down interactions and associated cascades as well as non-feeding related processes of the European bison (i.e. trophic rewilding (7)) (2).

In the Late Pliocene and Early Pleistocene the genus *Bison* appeared widely in the temperate regions of Asia and Europe giving origin to different forms of bison including *Bison priscus*. *Bison priscus* was present in Western Europe until the Late Pleistocene (10-12 kya) and is thought to have been replaced there by the modern European bison, *Bison bonasus,* which emerged in southern Caucasus after the Last Glacial Maximum (22kya) (8) and inhabited northern Germany and southern Scandinavia approximately 10kya (9). By mid-Holocene (5kya) *Bison bonasus* had disappeared from Western Europe, and later it also disappeared from Central and Eastern Europe (10). The decline of *Bison bonasus* is thought to be caused by increasing intensification of agriculture leading to habitat destruction and fragmentation forcing the escaping bison into the forest of Eastern Europe (8). By the late 1800s, there were only two European bison populations left, and the last wild bison died of disease in 1927 (10). After World War II, a breeding and reintroduction program was initiated based on 12 individuals out of the 54 individuals surviving in zoos across Europe (8). Reintroductions have until recently primarily occurred in forested areas (4) as European bison have been thought to be a forest specialist originating from closed forest habitats. However, recently more studies report that bison originally roamed in mosaic landscapes foraging on grassland plant species (9, 11–14).

The extensive reintroduction work has so far paid off, as there now are 1647 bison in captivity, 400 semi-free bison and 4009 free-living, mainly located in Poland, Belarus, Russia, and Caucasus (15) and the IUCN red list category has been moved from ‘endangered’ to ‘vulnerable’ in 2008 (16). However, challenges still remain as the free-living bison are distributed on 40 small and rather isolated populations with no or little genetic exchange (15). None of these wild populations are considered to be self-sustaining (17).

Despite the many reintroductions of European bison across Europe, there is still no clear roadmap for how to successfully reintroduce bison in highly anthropogenic landscapes (3) where habitats often are size-restricted. Consequently, it is even more important to match the habitat requirements of bison to the reintroduction site in order for it to be successful. Reintroduction efforts should, therefore, undergo evaluation to help fill in the many knowledge gaps e.g. in which habitat types do bison seem to thrive and under which management protocols do they display their ecological functions optimally.

In Denmark two small semi-wild herds have been established; in the riparian meadows of Gudenå river close to the city Randers in Jutland since 2010 (5) and since 2012 in the forests of Almindingen on Bornholm Island (18). The habitat use of two individuals of the herd consisting of 11 individuals on Bornholm has been evaluated using GPS (19). This study will be the first to study the case in the peri-urban meadow in Jutland.

In this study, we provide an assessment of the daily and seasonal behaviour of the bison and compare it to that of free-ranging herds. We also investigate the bison herd´s habitat use and selection by linking the occurrence of the bison herd to environmental parameters such as habitat characteristics, plant community traits, topography, and management area (release area with supplementary feeding vs. seminatural meadow area) to better inform future reintroductions of European bison and rewilding-inspired efforts in terms of assessing suitable habitat and management interventions.

## 2. Material and methods

### 2.1 Study area

The study was conducted in a 40 ha large bison enclosure located along the river Gudenå, in the Eastern part of Jutland, western Denmark (Fig 1A). The area mostly consists of wet meadows (79.7 %) partly susceptible to flooding, small deciduous forest areas (10.3 %), and previously cultivated grassland (5.2 %) (Table 1). The bison enclosure consists of two areas (Fig 1B), a release area (Fig 1D-E) and a meadow area (Fig 1C), between which the bison have free access most of the year. Formerly, the release area had been used for hay-harvest followed by cattle grazing. The release area contains a permanent stable, hay rack, and a water container (Fig 1D-E). The release area is located 0-3 m above sea level (20) and is still being drained by use of drainpipes and ditches, though less effectively as during former land use practices. Parts of the meadow area have undergone a cutting regime in the fall of 2013 and 2014 as these parts are enrolled in the Danish Common Agricultural Policy payment scheme for management of grasslands and nature areas. Roe deer (*Capreolus capreolus)* have been spotted inside the bison enclosure, as well as hares (*Lepus europaeus*), stoat (*Mustela erminea*) and foxes (*Vulpes vulpes*).

**Fig 1:**
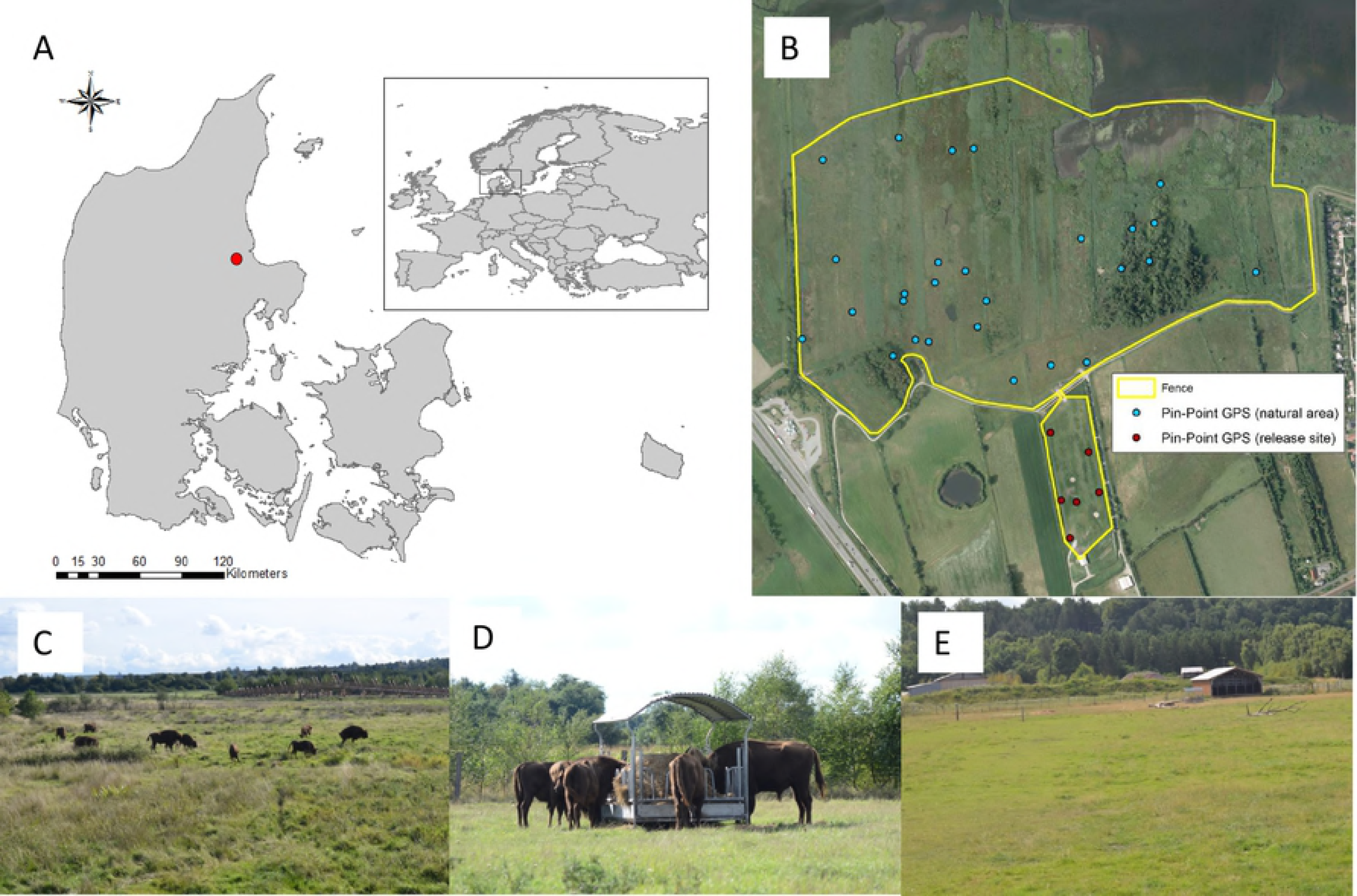
Overview of and photos of the study area. (A)The big map shows Denmark with the study location marked with a red dot and the inserted map shows the location of Denmark in Europe. (B) Ortho photo of the bison enclosure (yellow line) showing the GPS points used for the vegetation analysis within the meadow (blue points) and within the release area (red points). (C) Photo of the bison herd feeding in the meadow. (D) Bison herd feeding at the hay rack in the release area. (E) Photo of the release area with the barn in the far end.

**Table 1:**
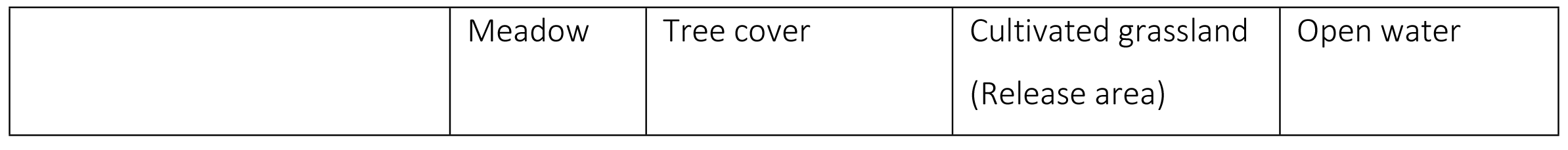
Time spent by the bison herd in each type of habitat compared to the availability of the habitat types.

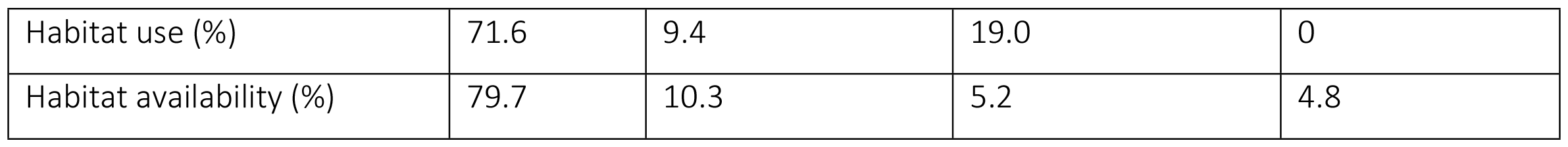

### 2.2 Study animals

Here we study 11 European bison of the lowland line (*Bison b. bonasus*) (1 bulls, 4 cows, and 6 female juveniles) that arrived at the bison enclosure in the summer of 2012. When the bison first arrived, they were initially kept in the release area, which is positioned furthest away from the open water and highest above sea level. During winter, they receive supplementary feeding, provided in the release area. Supplementary fodder is also available during other times of the year as a means to apply veterinary treatment to the bison and to fulfil Danish legal animal ethics e.g. animals should stay in good body condition year-round. During the study period, bison were observed feeding in the area with supplementary fodder more often than in the rest of the release area, indicating that bison exploited this supplied resource (mean number of feeding observation in grid cells with supplementary fodder 4.2).

### 2.3 Data collection of bison behaviour, habitat use and plant species

Observations on behaviour and position were made from April 29 to September 27, 2013. During this period, observations were made 2-3 days a week on average, with a total of 44 observation periods. Each observation period lasted 7-9 hours, where behaviour and position were logged every 15 minutes. Temperature, time and weather conditions were also logged, as well as notes about unusual or relevant events. Observation periods were either ranging from dawn until noon, or from noon until dusk, where the herd would lay down for the night. In total, 22 cycles from dawn to dusk were obtained, summing up to 326 observation hours in total and resulting in 1380 observation data points. The whole observational period (April to September) was divided into three seasons (spring, summer, and fall) based on the average daily temperature (Fig S1), with the summer period being significantly warmer than the spring and the fall period (Dunn´s test: p=0.0001 and p=0.0004, respectively). Observations were made from outside of the enclosure and binoculars were used if necessary in order to categorize the behaviour of the bison. The behaviour observed was divided into four categories (feeding, resting, moving, and other), inspired by previous bison studies (21, 22) (Table 2). For analysis of seasonal behaviour the type of behaviour was designated to the entire herd based on the type of behaviour the majority of the herd showed at the logging time. This observational study was approved by the organization responsible for the animals, Randers Regnskov.

To map the vegetation in the bison enclosure, 36 vegetation plots were randomly positioned based on random geographic coordinates assigned in R (23) (Fig 1B). Every plot location was found using a Trimble Juno SB GPS, however, three plots were positioned in open water and therefore discarded. In each plot, which consisted of a circle of 10 meters in diameter, plant species were identified on a presence/absence level. Access to the area and execution of floristic analysis was done in accordance with and approved by the organizations involved, Randers Regnskov and Aage V. Jensens Foundation.

**Table 2:** Description of the four different types of behaviour monitored while observing the bison herd.

| Behaviour type | Description |
| --- | --- |
| Feeding | The animal is standing still or taking slow steps without lifting its head. |
| Resting | The animal is lying down on the ground ruminating or sleeping. |
| Moving | The animal is walking or running with its head lifted off the ground and not eating. |
| Other | The animal is standing still with its head lifted. Nuzzling or scratching, playing or fighting, urinating or defecating falls under this category. |

### 2.4 Analysis of behavioural patterns and habitat use

Data on behaviour and position were processed in ArcGIS 10.1 (24). The study area was divided into 20×20m grid cells. This grid size was chosen based on the observed space used by the 11 bison. The position and type of behaviour were summed for each grid cell across the whole study period to link the spatial habitat use to predictor variables. Type of behaviour was summed for each grid cell across each defined season to analyse behaviour across season. Time spent on feeding, resting and moving was tested against previous findings by Cabon-Rackzynska et al. (22) on free-living European bison in Bialowieza Forest. This study reported that bison on a daily basis forage 60.4 %, rest 31.9 %, and move 7.7 % during periods with no snow cover. The percentage of each habitat type in each grid was calculated. Spatial analysis was conducted in R (23). One grid cell was removed from analysis involving Frequency of occurrence as we considered it an outlier as bison often were observed here (99 times) due to this grid cell being the physical link between the two Management areas (release area vs. meadow area with tree patches) of the enclosure.

We tested the difference in Frequency of occurrence among habitat types (cultivated field, meadow, water, and tree covered areas) with a Pearson´s Chi-squared test and difference in Frequency of occurrence among habitats when accounting for habitat availability was also tested pairwise with a Pearson´s Chi-squared test. Mann Whitney t-tests were used to test if the behavioural pattern or Frequency of occurrence differed across Management area (release area vs. meadow area). Whether the behavioural pattern of the bison herd differed across season (spring, fall, and summer) was tested with a Kruskal Wallis test, and significant test results were followed by pairwise test using Dunn test using no p-value correction. Correlation between time spent on a certain behaviour (feeding or resting) and the daily average temperature was tested with a corrected Pearson´s correlation test accounting for temporal autocorrelation. All statistical tests were performed in R (23).

### 2.5 Analysis of habitat selection

We considered two response variables (Fig 2A-B); the frequency of occurrence of bison in each grid cell in the total enclosure and presence/ absence of bison in each grid cell in the total enclosure. Frequency of occurrence was used as a measure for how often do the bison herd occur in a certain grid cell. Presence/absence was used as a measure for whether or not the bison herd occur in a certain grid cell. We considered five predictor variables (Fig 2C-F, Table 3). Three describe the local environment: Elevation derived from the Digital Terrain Model (25), Tree cover, and Management area (release area vs. meadow area with tree patches) and two describe variation in the plant community; Forage quality for cattle (26) retrieved from BiolFlor (27) and Specific Leaf Area (SLA: leaf area per leaf dry mass) retrieved from LEDA traitbase (28) in order to explain the habitat use by the bison. Elevation was included to gain insight about the role of hydrology on habitat selection as the enclosure ranged from a submerged area close to the river to a dry field furthest away from the river. Tree cover was a relevant variable to include considering the ongoing debate about whether bison prefer open habitats, mosaic-landscapes or closed forests. Management area (release area with supplementary feeding vs. semi-natural meadow area with tree patches) was considered to assess the influence of the enclosure configuration and management protocols as this might affect their habitat selection and behaviour. Forage quality and SLA were included as it is likely that bison prefer areas where plants species provide high forage quality or digestibility, which has been found to increase with increased SLA (29) and might constitute a relatively easy measure predicting where bison impact should be expected.

SLA and Forage quality were interpolated to the whole study area based on presence/absence data on plant species obtained from the 33 vegetation plots in order to include these variables in the spatial models (Fig 2). We used kriging with the following co-variables; Elevation and Digital Object Model (a measure for vegetation structure), which was derived from Elevation and Digital Surface Model (25). Performance of co-kriging was tested using a training subset and a validation subset of vegetation plots and resulted in a prediction of 67.2% for SLA and of 85.1% for Forage quality. Co-kriging was performed using the function krige() in the R package gstat (30, 31) and based on exponential variogram shape.

The response variable Frequency of occurrence was log transformed to ensure normality of the residuals, and all grid cells with zero values (no observations of bison) were removed to avoid skewness and bias in the residuals. One grid cell was removed from analysis involving Frequency of occurrence as we considered it an outlier as bison often were observed here (99 times) due to this grid cell being the physical link between the two Management areas (release area vs. meadow area with tree patches) of the enclosure.

The relationship between the occurrence frequency (the number of times the bison herd occurred in the same grid cell) and Elevation, Tree cover, Management area, Forage quality and SLA was tested with simple linear regressions for continuous explanatory variables and with Pearson product moment correlation for categorical explanatory variables.

Multicollinearity among predictor variables was tested using Spearman’s rank correlations for variables with a non-linear relationship and Pearson´s correlation between variables with a linear relationship. Multicollinearity was considered a problem among predictor variables when correlations rose above 60%, and therefore Management area and Forage quality were excluded from the habitat selection models as they were highly correlated with Elevation (See Table S1). Elevation was not discarded as this predictor variable was of highest resolution.

Spatial autocorrelation was tested by evaluating Moran´s I for the lower distance classes of the residuals and considered to be negligible (see Fig S2). Linear regression models were conducted to test if the predictor variables (Elevation, Tree cover, and SLA) influenced how often the bison herd occurred in a given grid cell. Multiple logistic regression models were performed to test if the predictor variables (Elevation, Tree cover, and SLA) had an effect on whether the bison occurred in the grid cell or not (presence/ absence). All possible combinations of models (eight models for both linear and for logistic models) were fit and ranked according to their relative weight of evidence (using Akaike weights, w_i_) calculated from Akaike’s Information Criterion corrected for small sample size (AIC_c_). Models were assessed according to the strongest support (w_i_) and explanatory power (33). To evaluate the explanatory power of each individual variable, we calculated model-averaged coefficient estimates (parameter estimates weighted according to support across all eight models) and relative variable importance (RVI, the sum of Akaike weights for each model in which the explanatory variable occur) (33). Spatial model selection and model averaging were conducted using R package MuMIn (34).

We also looked into an old study assessing the natural food preferences on European bison in Poland (32) to investigate if the results of this study supported a relationship between feeding preference and SLA. We therefore assigned SLA values to all the plant species for which Borowski and Kossak (32) had measured abundance coefficients (D, after German “Deckungswert”) and contacts between bison and plant (n) (bites) and tested the relationship between food preference and SLA (n/D~SLA) with a linear regression.

### 2.6 Ethics statement

The animals were kept in a 40 ha enclosure privately owned by the NGO Aage V. Jensen Nature Foundation. The zoo Randers rainforest purchased six animals from UNK at Bialowieza National Park and five animals were donated to the zoo Randers Rainforest from another Danish zoo Ree Park – Ebeltoft Safari. The zoo Randers Rainforest was responsible for animal handling, control, release and zookeepers supervised the animals daily. The physical body condition of the animals was regularly checked by the supervising veterinarian of the zoo Randers Rainforest. This study did not imply animal sampling of protected animals.

**Fig 2:**
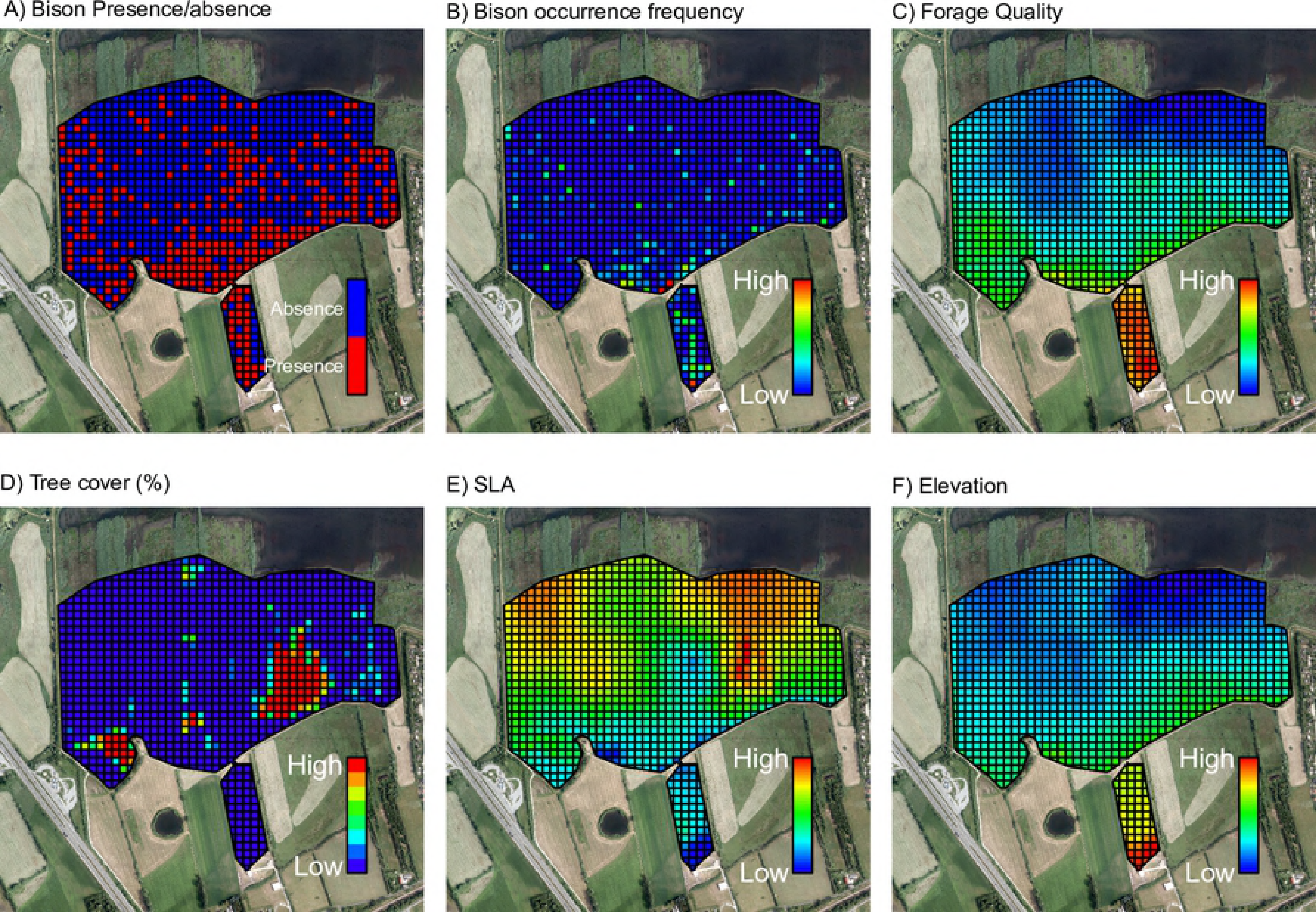
Response and predictor variables used for the spatial linear and logistic regression. Response variables: (A) Response variable: Presence/ absence and (B) frequency of occurrence of the bison herd summed across the whole observation period. Predictor variables: (C) Forage quality, (D) Tree cover, (E) SLA, and (F) Elevation.

**Table 3:**
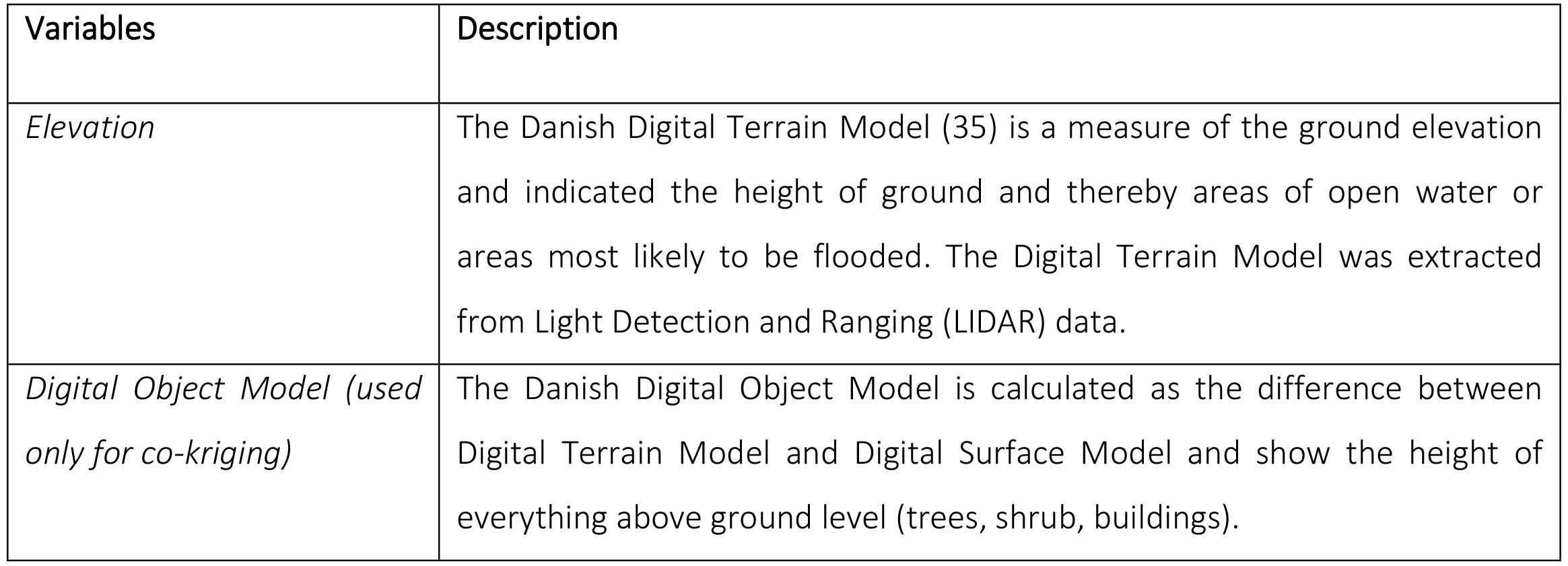
Description of the predictor variables used in the spatial linear and logistic regression.

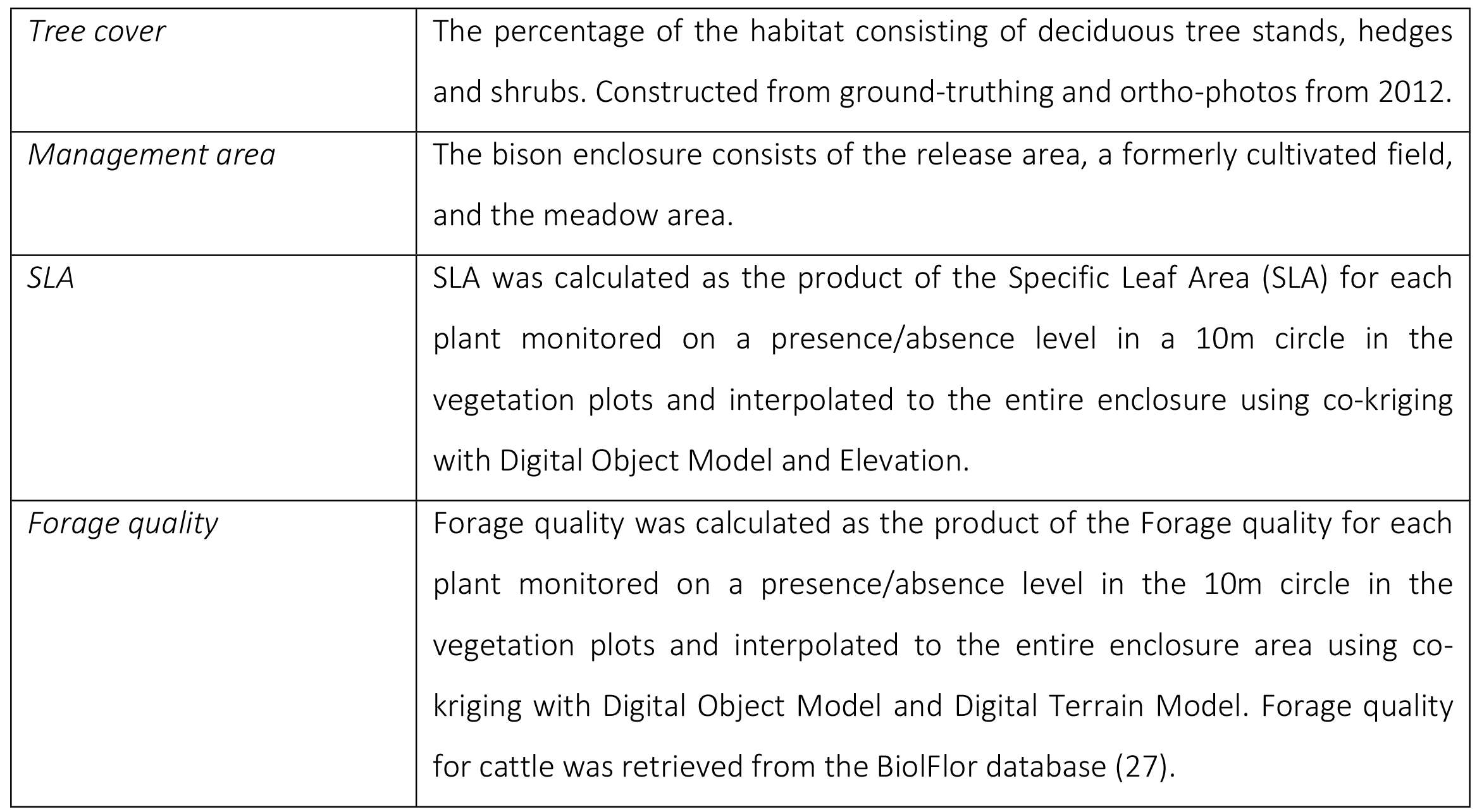

## 3. Results

### 3.1 Behavioural patterns

The diurnal behaviour pattern showed that the bison herd had three major feeding bouts a day (Fig 3A), with intervening resting bouts. Over the entire growth season, the bison herd spent on average 59.4 % on feeding, 29.5 % on resting, 3.3 % on moving, and 7.8 % on other activities (Fig 3B). This activity budget was consistent with previous findings by Cabon-Rackzynska et al. (22) (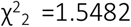, p=0.461). Time spent on resting was more or less unchanged over the three seasons (spring, summer, and fall) (Fig 3B), though time spent on resting was significantly correlated with the daily temperature (r= 0.38, F_40.58_=7.04, p=0.0113, corrected Pearson correlation for temporal autocorrelation) (Fig 4A). There was a non-significant tendency for the bison herd to spend more time on feeding as the growth season progressed (H_2_=5.48, p=0.0647, Kruskal Wallis Rank Sum test) (Fig 3B). Time spent on feeding and daily temperature were not correlated (r=0.03, F_23.7758_=0.0257, p=0.87) (Fig 4B). Time spent on moving and other behaviour types showed a significant change across season with the spring period showing significantly higher moving activity than the summer period (p=0.0020, Dunn′s test) as well as a higher level of other behaviour (e.g. scratching and playing) during spring time than the summer and fall period (p<0.0001 and p=0.0217, respectively, Dunn´s test) (Fig 3B).

**Fig 3:**
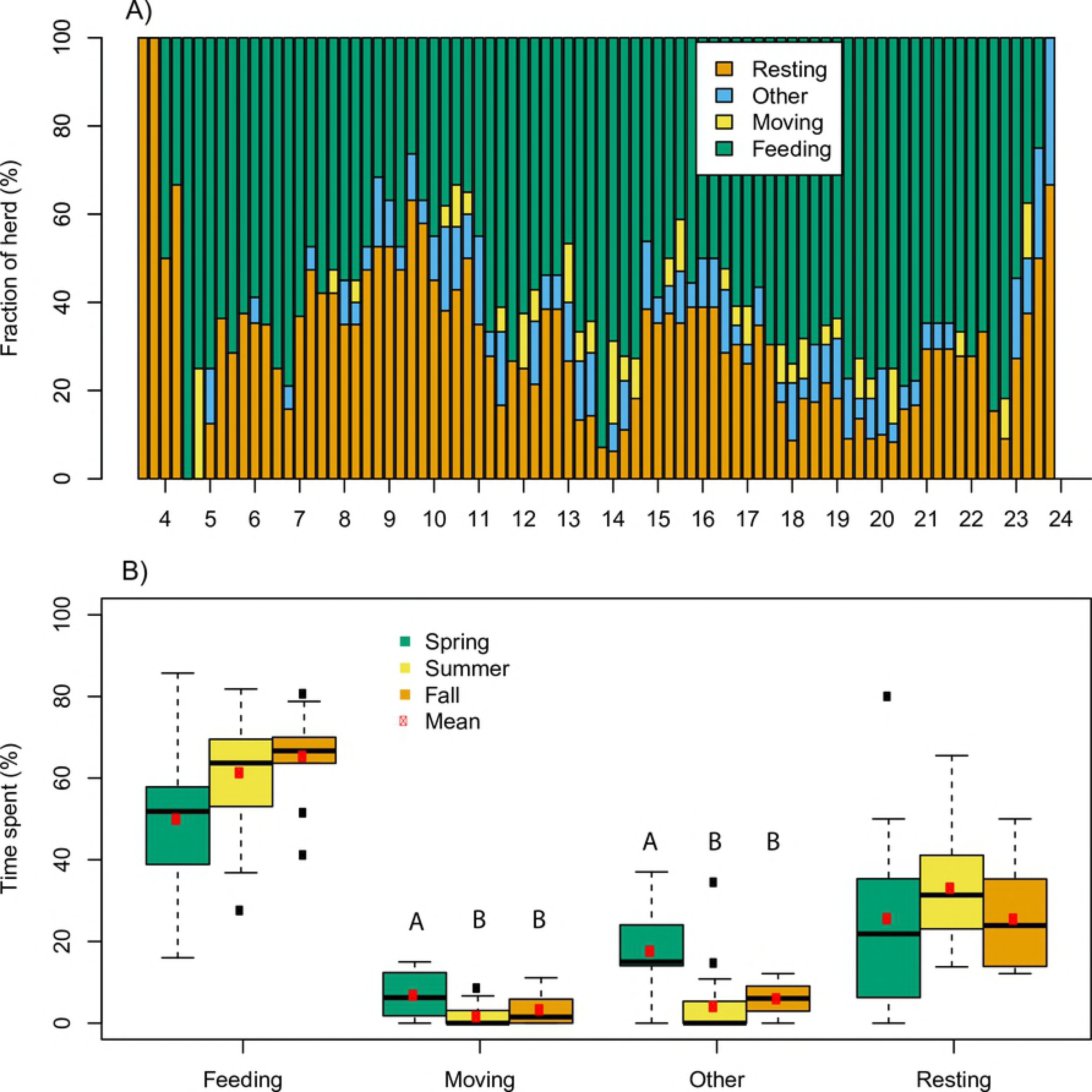
Daily and seasonal behavioural patterns. Average percentage of individuals in the bison herd resting, feeding, moving or expressing other types of behaviour during a day throughout the observation periodl (A). Time spent on a specific type of behaviour across the seasons (spring, summer, and fall) (B). Different letters are attributed to seasons that vary significantly from other seasons within the same type of behaviour.

**Fig 4:**
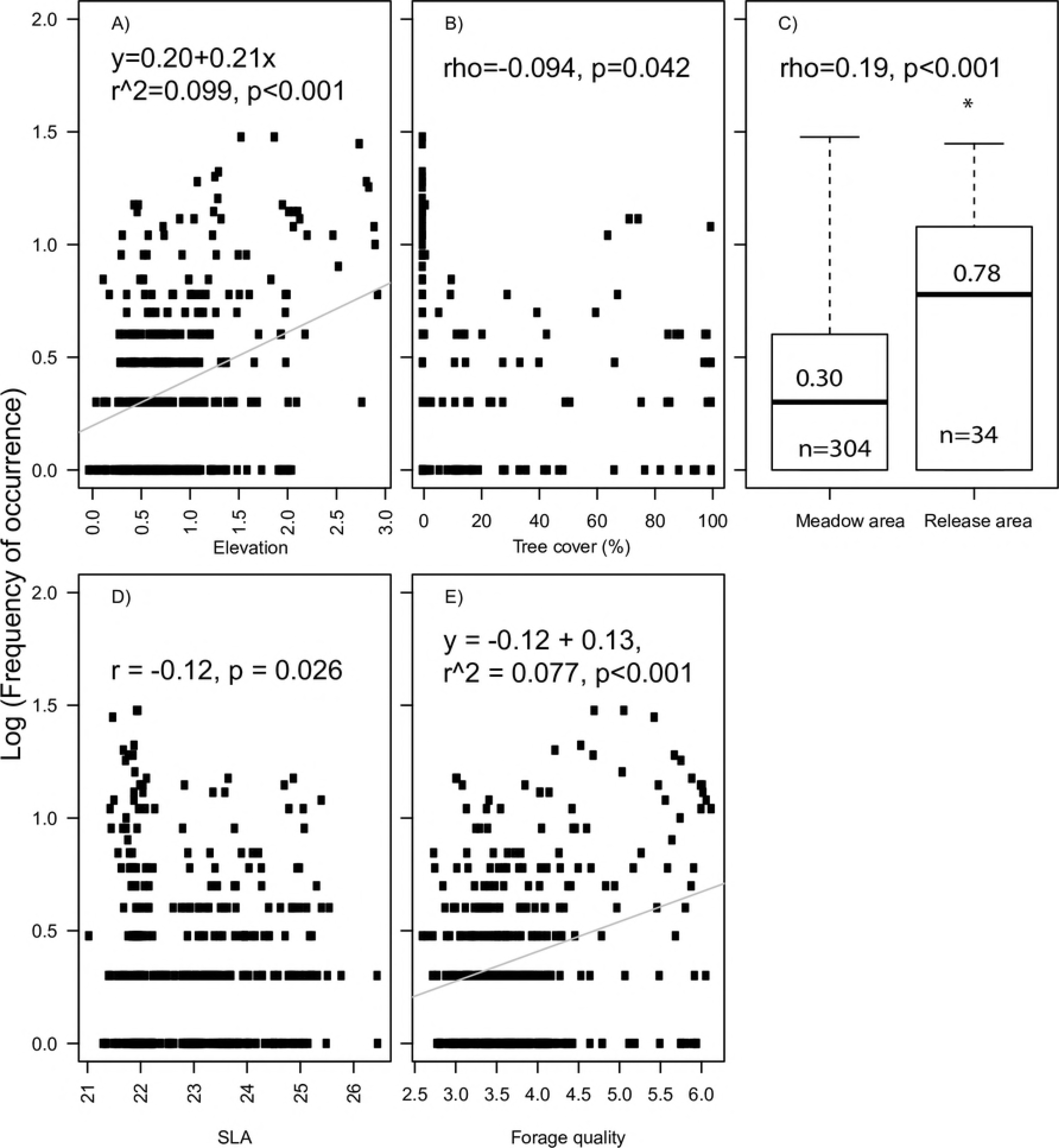
Correlation between temperature and time spent resting or feeding. Correlation between mean daily temperature and time spent on (A) resting or on (B) feeding across the observation period. Statistical results from Pearson´s correlation corrected for temporal correlation are shown.

### 3.2 Habitat use

Overall, the bison herd spent significantly more time in the open meadow than in any of the other types of habitat (the cultivated release area, the forest patches, and open water (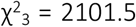, p<0.001), but not as much time as predicted from the availability of the open meadow area (Table 1). Bison spent almost the quadruple amount of time in the release area than expected from its´ availability (Table 1), which was significantly more than time spent in other habitats; meadow, areas with tree cover, and significantly more than in the other management area (meadow with patches of open water and trees) (Table 4). There was no difference in habitat use of meadow habitat and areas with tree cover or between open areas (meadow combined with release area) and areas with tree cover (Table 4).

Log frequency of occurrence was significantly correlated with management type (Spearman´s rank correlation: rho= 0.19, S=5211000, p<0.001) and the number of resting and feeding observations per grid cell in the release area was significantly higher than in the meadow area (Wilcoxon w=3534, p<0.0001 and Wilcoxon w=3678.5, p<0.005) (Fig 6). We found no strong relationship between log frequency of occurrence and any explanatory variables, though there were moderate to weak relations to Elevation (r^2^=0.099, p<0.001), SLA (r=−0.12, p=0.026) and Forage quality (r^2^=0.077, p<0.001) (Fig 5).

**Fig 5:**
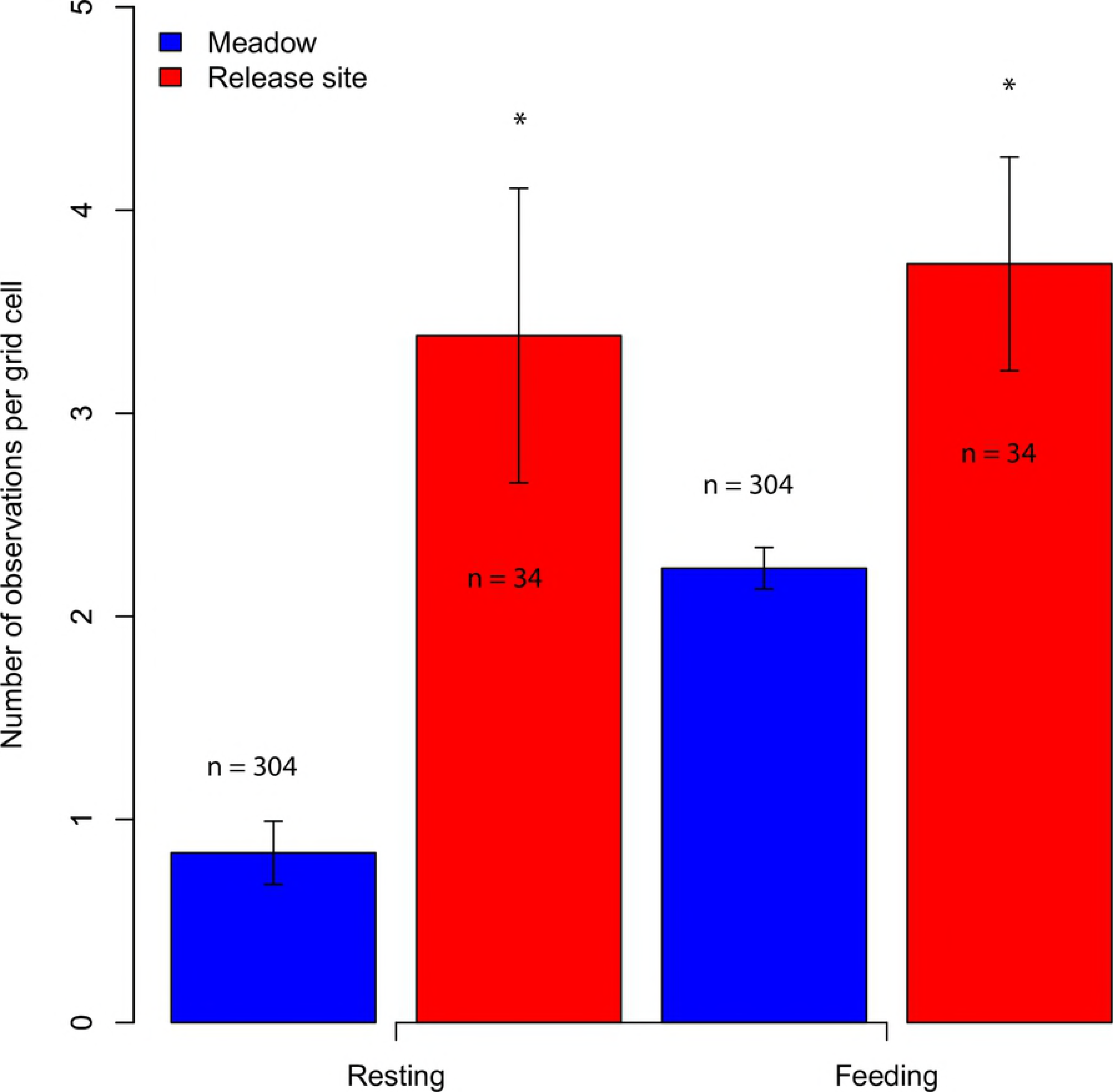
Correlations between Frequency of occurrence and predictor variables. Correlations between log frequency of occurrence and (A) Elevation, (B) Tree cover, (C) Management area, (D) SLA, and (E) Forage quality). Statistical results from Pearson´s correlation or Spearman´s rank correlation is shown for continuous variables and ordinal variables. An asterisk indicates a significant difference among Management areas (C).

**Fig 6:**
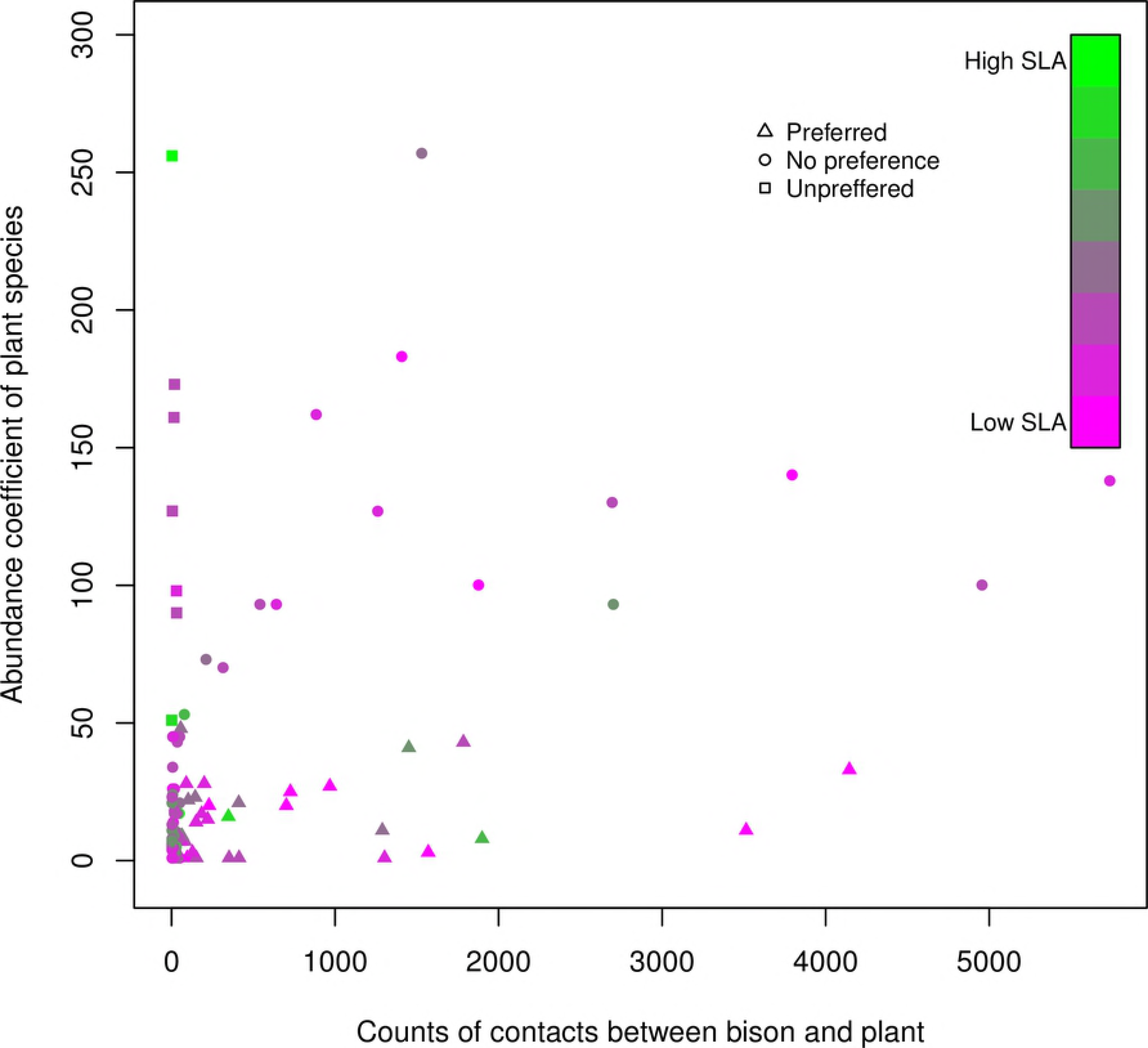
Time spent feeding and resting across management area. Number of feeding events or resting events of the bison herd observed in the meadow area or in the release area. n value indicates number of grid cell per mangement area. An asterisk indicates a significant difference in number of observations between management area.

**Table 4:**
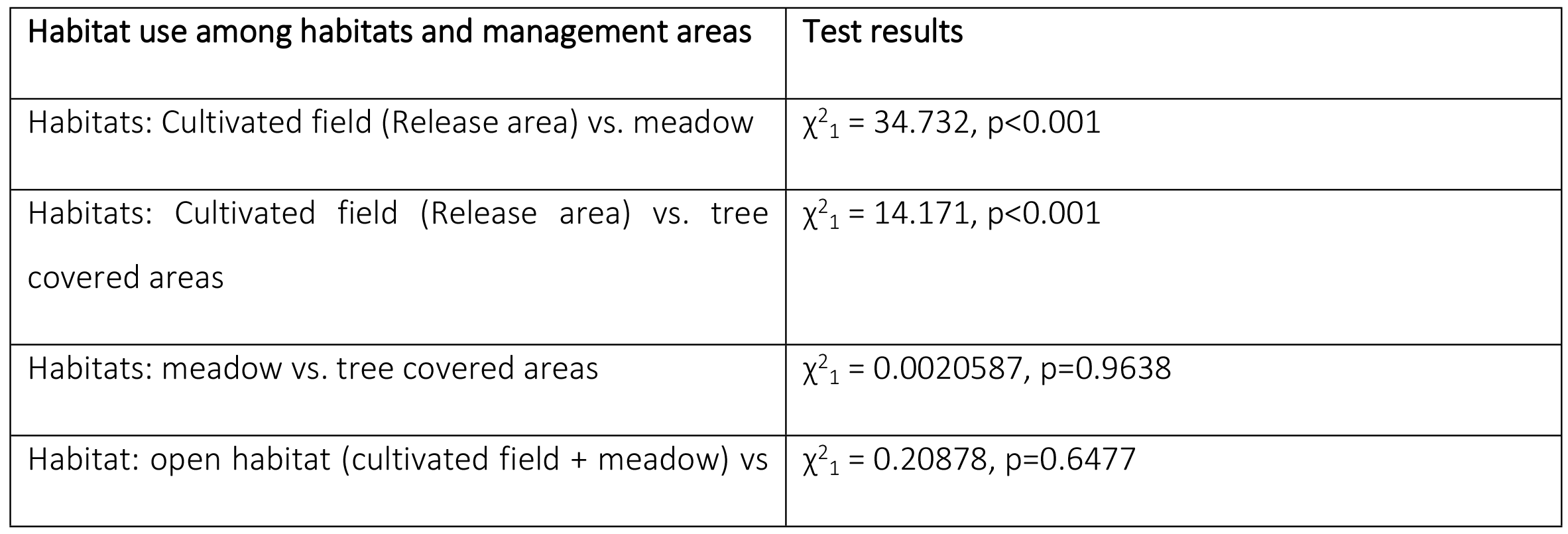
Statistical results of chi-square test of habitat use of different habitats and managements areas when accounting for their habitat availability.

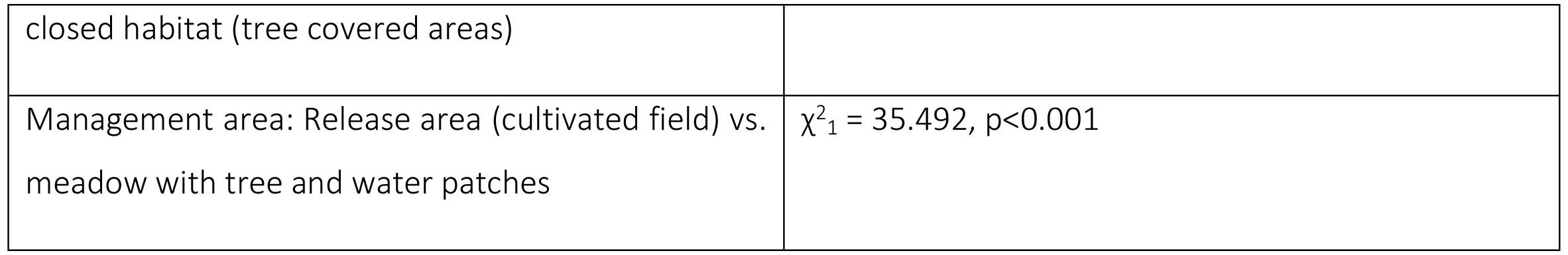

### 3.3 Multiple linear and logistic models of habitat selection

None of the models considered showed problematic levels of spatial autocorrelation. Therefore, spatial linear models were performed with Ordinary Least Square and spatial logistic models were performed with Generalized Linear Model (Fig S2). The three best linear regression models (ΔAIC_c_ < 2) all have fairly low w_i_ values and low adjusted r^2^ values (Table 5), indicating that the models neither are notably better than the worst models nor have great explanatory support. These linear regression models indicate that Elevation is the most important predictor as it appears in all the best three models and has the highest effect in the linear regression models, while Tree cover and SLA both have minimal effect (Table 6). Elevation had positive coefficients (Table 5), suggesting that bison preferred staying in areas with higher elevation, which were also drier than the area with lower elevation.

For the logistic regression models, the best models seemed to have much higher explanatory support than the second best model (w_i_ = 0.85 vs. w_i_ = 0.15, respectively), though both had low predictive power (McFadden’s r^2^=0.0830 and 0.0790). All explanatory variables were included in the best model, whereas the second best model did not include Elevation. Elevation and Tree cover had positive coefficients, whereas SLA had negative coefficients (Table 7). SLA had the highest effect for the logistic regression models, and Tree cover and Elevation had moderate effects (Table 6). The logistic model indicates that the bison herd more likely occurred in areas with higher elevation, like the linear regression model, but also in areas with lower SLA-values and more tree cover. Overall, these models suggest that the bison herd visited high-lying, drier areas more often than low-lying areas, wet areas, and that the bison herd less likely visited areas with high SLA-values at all.

Assigning SLA values to the bison food preference data obtained by Borowski and Kossak (32) allowed us to test the relationship between food preference and SLA (n/D~SLA) (Fig 7). The linear regression had very low explanatory power and SLA was a poor predictor of food preference (adjusted r^2^=0.00082, p=0.30).

**Fig 7:**
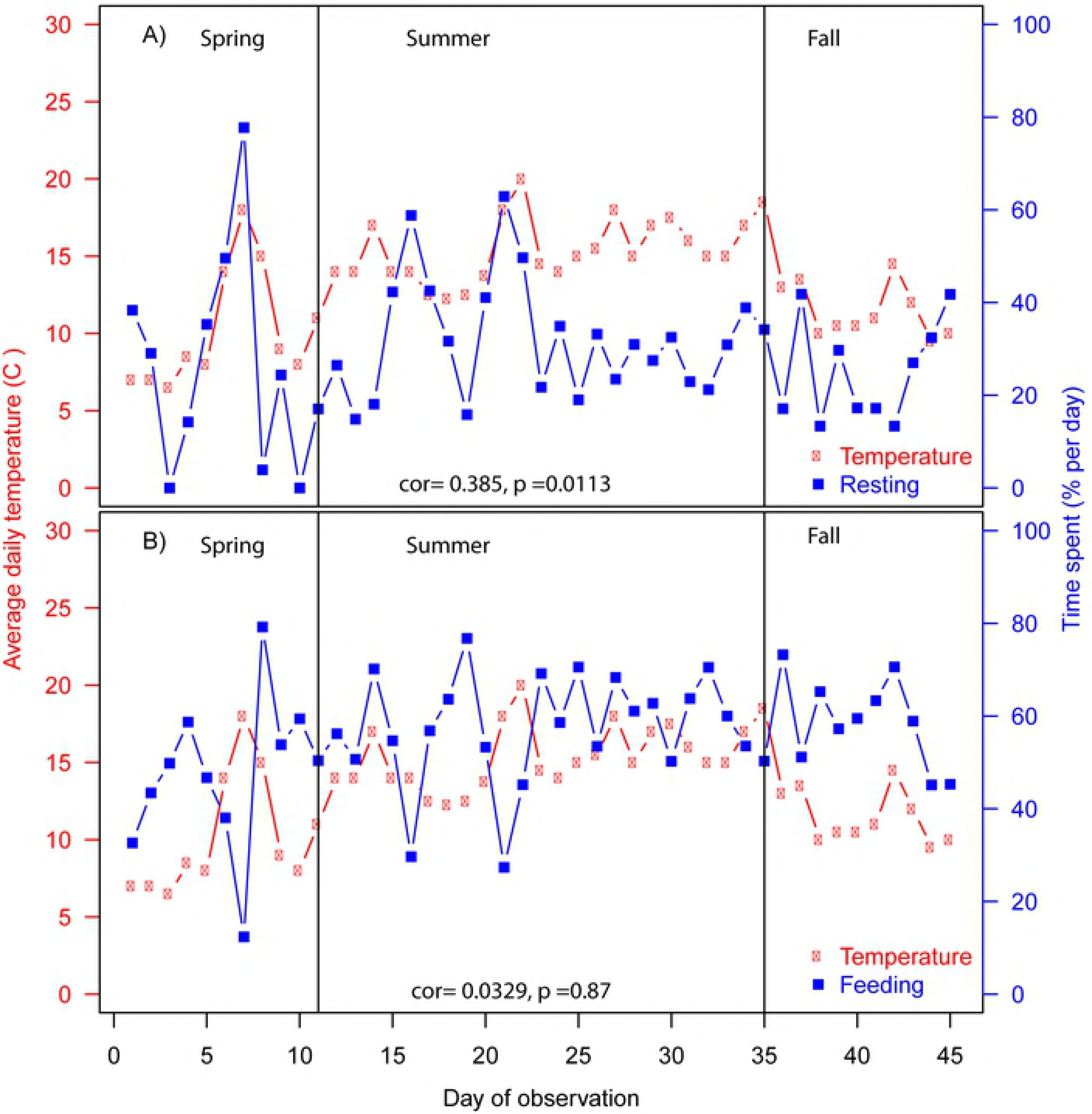
Relation between plant preference and SLA-values. Correlation between plant-bison contacts, availability of plant species, and SLA-values of plant species. **Unpreferred plant species** (rectangle) is characterized by having contacts with a bison less than expected by chance (an abundance coefficient of 50 or above while having less than 50 contacts with a bison), **Preferred plant species** (triangle) is characterized by having contacts with a bison more often than expected by chance (an abundance coefficient less than 50 while 50 or more contacts with a bison), and **plant species with no preference** (circle) is characterized by having as many contacts with a bison as expected by chance (an abundance coefficient above 50 while having more than 50 contacts with a bison or an abundance coefficient equal to 50 or lower while having 50 or less contacts with a bison). Figure adapted from Borowski and Kossak,1972.

**Table 5:** Three best spatial linear models on log frequency of occurrence (ΔAIC_c_ <2) selected from eight tested models and ranked by AIC_c_.

| Model coefficients (n=1095) | | | $\Delta\text{AIC}_c$ | $w_i$ | Adjusted $r^2$ |
| --- | --- | --- | --- | --- | --- |
| Tree cover | Elevation | SLA |  |  |  |
| -0.08 | 0.37 |  | 0 | 0.317 | 0.141 |
|  | 0.37 |  | 0.27 | 0.277 | 0.138 |
| -0.10 | 0.42 | 0.08 | 1.73 | 0.273 | 0.143 |

$\text{AIC}_c$ (corrected Akaike Independent Criterion); $w_i$ , Akaike weights.

**Table 6:** Summary of results after full model-averaging: effects of each explanatory variable on Frequency of occurrence (linear regression) and presence/absence (logistic regression) of bison.

| Predictors | Linear regression |  |  |  | Logistic regression |  |  |
| --- | --- | --- | --- | --- | --- | --- | --- |
|  | Estimate | Adj. SE | CI 95% | RVI | Estimate | Adj. SE | CI 95% |
| Elevation | 0.391 | 0.0587 | 0.275 –<br>0.506 | 1 | 0.435 | 0.227 | -0.083 –<br>0.807 |
| SLA | 0.0291 | 0.0545 | -0.0777 –<br>0.136 | 0.59 | -1.28 | 0.221 | -1.71 –<br>-0.844 |
| Tree cover | -0.0514 | 0.0592 | -0.167 –<br>0.0647 | 0.41 | 0.630 | 0.159 | 0.318 –<br>0.941 |

Adj. SE, Adjusted standard error; CI 95%, 95% confidence interval; RVI, relative variable importance.

**Table 7:** The best spatial logistic model on presence/ absence occurrence (ΔAIC_c_ <2) selected from 8 tested models and ranked by AIC_c_.

| Model coefficients (sample size: n=338) | | | $\Delta AIC_c$ | $w_i$ | McFaddens $r^2$ |
| --- | --- | --- | --- | --- | --- |
| Tree cover | Elevation | SLA |  |  |  |
| 0.61 | 0.42 | -1.2 | 0 | 0.85 | 0.0830 |
| 0.72 |  | -1.5 | 3.47 | 0.15 | 0.0790 |

$AIC_c$ , (corrected Akaike Independent Criterion); $w_i$ , Akaike weights.

## 4. Discussion

In this study, we wished to assess the daily and seasonal behaviour of bison and compare with previous findings in Bialowieza Forest. The bison herd had three major feeding bouts a day (Fig 3A) and on average spent 59.4 % on feeding, 29.5 % on resting, and 3.3 % on moving across the growth season (Fig 3B), which was consistent with previous findings on bison in Bialowieza Forest (22). We also wanted to investigate if the bison herd´s habitat use and selection were linked to environmental parameters such as habitat characteristics, plant community traits, topography, and management area (release area and semi-natural meadow area). We found that bison used the release area more than expected from availability, but did not favour open habitat over forested habitat when accounting for habitat availability. We also found that they preferred drier high-lying areas and areas with low SLA values over wetter low-lying areas and areas with higher SLA values.

### 4.1 Behavioural patterns

We found that the herd of European bison had three major feeding bouts a day on average; the first around dawn, the second around noon and the third and final around dusk. Previous studies from Bialowieza Forest report four feeding bouts a day in periods without snow cover (22) and in periods with snow cover two feeding bouts have been reported (21). Over the entire growth season the bison herd spent on average 59.4 % of their time feeding, 29.5 % resting, and 3.3 % moving, which was consistent with the findings on behaviour of bison in Bialowieza Forest during periods with no snow cover (22). The bison herd in Vorup meadows overall seem to behave like free-roaming bison observed in large forested areas less human-modified, despite being confined to a heavily human-modified meadow enclosure. This is an important finding as this indicates that rewilding with European bison in anthropogenic environments can support thriving animals expressing what we, so far, believe is natural behaviour in the wild. Knowing this also enables us to transfer knowledge from one reintroduction site to another more confidently.

Bison reintroduced in the Alps in France with access to supplementary fodder spent more than 40% of the time feeding and approximately 50% resting independent of snow cover (3). In contrast, Cabon-Rackzynska (21) found that bison in Bialowieza forest, also with access to supplementary fodder, spent less time of the day feeding (30% vs. 60%) and correspondingly spending more time resting in periods with snow cover. Resting more during cold periods can be related to shifts in temperature and weather conditions, which appear to cause the European bison to behave more docily (22). Our observations also support that shifts to higher temperature increase the amount of time spent on resting (Fig 4A).

### 4.2 Habitat use

Overall the bison herd preferred open areas over patches with tree cover, and the herd spent most of its time in the large meadow area (~72 %) (Table 1). Nevertheless, the herd actually spent less time in the meadow area than expected from its area coverage (Table 1), coupled to it spending more time (19%) in the release area than expected from availability (~5%) (Fig 5C and Table 1). It is unclear whether the preference for the release area is linked to its inherent ethological value (i.e. the possible status of the release area as their “home” might influence their behaviour accordingly), or to the rather high forage quality of the release area (Fig 2C) due to former grass sowing management or the fact that the managers from time to time supplied fodder in a hay rack, or that the release site was placed at the driest part of the enclosure. It is reasonable to think that the preference for the release area is linked to feeding habits as the herd was observed significantly more times feeding in grid cells in the release area than in the meadow (Fig 6), however, the herd was also observed significantly more time resting in grid cells in the release area than in the meadow indicating that the release area has ethological value for the bison (Fig 6). Bison did not show any preference between open habitats (release area + meadow areas combined) and areas with tree cover when accounting for their habitat availability, and our results thereby do neither favour that bison is a forest species or an open habitat species.

A Danish study on the bison population at Bornholm, Denmark, has reported that the bison prefer uncultured cut coniferous forest over cultured coniferous forest and deciduous forest in their 200 ha enclosure consisting primarily of 70 % deciduous and coniferous forest and 30 % meadow (19). European bison released to a former commercial plantation, Roothaargebirge, in Germany have been reported to show habitat preference for areas with spruce, storm-damaged areas and grasslands and habitat avoidance to pine forest and beech forest (4). An early study on free-living European bison in the Bialowieza forest have reported that the bison show more or less equal preference for mixed coniferous forests and fresh deciduous forest (36), however, a more recent study shows that the habitat preference of both the Polish and Belarusian bison population has shifted towards deciduous forest and alderwood (37). This tendency has been confirmed recently in another study on the Polish population (38). Free-ranging bison in Lithuania has also been reported to feed more on grass than any other food source throughout the year indicating their habitat preference for open habitats (39). Cromsigt and colleagues (2) recently investigated the diet of European bison reintroduced to 220 Ha of coastal dune landscape including konik horses in Kransvlaak, The Netherlands, and found that bison predominantly fed on grasses but also included approximately 20% of woody plant matierial in their diet. Compared to the diet of Higland cattle, studied in a neighbooring area to Kransvlaak, bison diet differed as bison overall debarked more than cattle – especially during winter and spring (2). Some studies also document that bison often rest in forest patches, presumably to avoid disturbance (11, 40). Kerley and colleagues (11) have found that more than two-thirds of the free-ranging bison herds have in fact expanded their range, from originally being forest habitat, to also include open habitats. This, however, is not considered a result of the bison´s habitat preference, but as a result of a management problem as the open habitats bison adopt often are agricultural lands. Conspicuously, 65% of the free-ranging bison herd are also provided with supplementary fodder in order to reduce unintended impacts of bison herds (11). These varying results indicate that European bison can survive in different habitats including open and semi-open habitats. The refugee species concept have been proposed for European bison, which suggests that Pleistocene and early Holocene bison occupied open habitats to a larger degree than forest habitats (9, 11–14). Kerley and colleagues (11) have suggested that the overall larger habitat use of forest by bison observed today is a consequence of anthropogenic pressure, habitat change, and the fact that conservation efforts have worked within the “bison as a forest species” paradigm.

Schmitz and colleagues recently investigated the exploration behaviour of a European bison herd reintroduced to a mountainous area in Germany during the transition from an 89 Ha release area (occupied for three years) to the total enclosure of 4500 Ha. They found that the release area was fully or partially included in the 85 % Kernel density (i.e. used areas) by the herd or the bull in 15 out of 18 10-days periods during the first 6 months of exploration (4). Though the authors explicitly state that bison only used the release area occasionally, these results indicate to us that the exploration behaviour by bison is centred around the release area in the majority of the time during the first 6 months and that the release area plays a significant role in how new habitats are occupied. Evidence exists from other studies that European bison show higher habitat preference for areas provided with supplementary fodder than if no supplementary feeding takes place (41, 42). Recently Ramos and colleagues (3)showed that a herd of European bison reintroduced in the French Alps spent more than 50% of the time in areas with supplementary fodder, but when no provision of fodder occur bison spent significantly less time by the food racks and exploited other areas to a higher degree (3). Influence of access to supplementary fodder on bison winter diet was recently investigated using DNA-barcoding by Kowalczyk and colleagues in Bialowieza Forest. Kowalczyk et al. (43) reported for bison occurring in forest habitats that woody material made up 16% of the diet of bison intensively fed with supplementary fodder, whereas the diet of non-fed bison was composed of 65% woody material (43).

When bison graze and spend time at a food rack they cannot exert their ecological function in other areas at the same time. This means that the ecological impact by bison is reduced not only in terms of grazing pressure but also in terms e.g. browsing, debarking (2, 43), trampling, distributing autochtonous nutrients, and seed dispersal, presumably lowering the potential of the bison to affect plant diversity, structure and composition (44), and thereby species communities dependent on these e.g. arthropods, amphibians, reptiles, birds and mammals (45). This might compromise or oppose conservation goals of trophic rewilding initiatives focusing on ecological restoration based on restoring trophic top-down interactions and associated cascades as well as non-feeding related processes of the European bison (7).

Reintroductions are often conducted in two steps; releasing the bison into to a smaller release area e.g. with supplementary fodder initially or continually, followed by allowing animals access to an adjoining, larger, and often more natural enclosure area (4). Based on the results presented in this study and other recent studies of bison reintroductions (3, 4, 43) we advocate that constellation of release area and final reintroduction area should be thoroughly considered along with how management protocols (e.g. supplementary fodder) might affect the habitat use by the animals and thereby conservation goals – particularly when these are framed within a trophic rewilding context.

### 4.3 Effect of environmental drivers on habitat selection

All explanatory variables (Elevation, Management area, SLA, and Forage quality) except from Tree cover showed significant correlations with Frequency of occurrence (Fig 5). These results indicate that the bison herd more frequently occurs on higher and drier areas with high Forage quality than on lower and wetter areas. The linear regression models showed that Elevation had the highest effect (Table 7).

Elevation is for this area related to wetness of the soil, as the enclosure is placed on the Gudenå River bank, with terrain rising up away from the water. The area is in the lower parts susceptible to flooding, which could be the reason why the bison herd seems to prefer higher grounds. In this study we did not observe the bison herd occupying the open water patches one single time; however, during the floristics analysis we saw obvious bite marks in areas partly submerged and bison have also been observed in water-filled ditches. In a global study, Olff et al. (46) linked plant-available nutrients and soil moisture to large herbivore densities and suggested based on this that large herbivores prefer drier areas as forage quality of plants are higher due to higher availability of soil nutrients. van Vuure has pointed out that bison´s preference for drier areas is reflected in the distribution of fossils, where only a few findings occur in river sediments (47). Unfortunately, in this study, we could not investigate the spatial relationship between habitat use and Forage quality as Forage quality was excluded from the spatial analysis due to multicollinearity between the Elevation and the Forage quality.

In the logistic regression models, SLA had a higher effect than Elevation (Table 7), suggesting that SLA was a better predictor of presence/ absence occurrence than Elevation and that SLA values often were low where the bison herd occurred. Whether bison actually avoided areas with high SLA values or not, is hard to deduct as Elevation and SLA were 59% correlated (See Table s1), just below the threshold for exclusion based on multicollinearity, and therefore the signals are hard to distinguish. Several of the herbaceous plants that European bison is reported to prefer (e.g. *Urtica diodica, Cirsium oleracium, Plantago sp, Filipendula ulmaria*) (48) were available in the enclosure, and in several vegetation plots, we observed clear bite marks on these species. Plants with high SLA values observed in the enclosure included e.g. *Impatiens capensis* (SLA 54.3 mm^2^/mg), *Urtica dioca* (SLA 31.6 mm^2^/mg), *Alisma plantago-aquatica* (SLA 29.7 mm^2^/mg), *Cirsium oleracum* (SLA 25.7 mm^2^/mg), and several grasses (SLA values from 18-35 mm^2^/mg). Some of the tree species (e.g. *Betula pubescens, Alnus glutinosa, Salix* sp.) reported as preferred (48) were also observed in the enclosure and showed clear signs of debarking. These species had lower SLA values than many herbaceous plants. One might think that plant species with high SLA values might be attractive for foraging bison due to their seemingly high palatability, but in this study, the logistic models indicated a negative relationship between occurrence and SLA. Testing the relationship between SLA and food preference for plant species (n/D) obtained by Borowski and Kossak (32) studying bison in Bialowieza Forest showed that SLA had very low explanatory power for food preference (adjusted r^2^=0.00082, p=0.30) (Fig 7). These results suggest that SLA is a poor predictor for the habitat selection of bison.

Our habitat selection results indicate that the environmental parameters underlying the bison herd´s habitat preference vary in terms of whether the bison herd chooses to use the area at all or choose to visit the area multiple times. There were three best models for Frequency of occurrence, while just one best model for presence/absence, this might suggest a rather different composition of areas chosen versus not chosen, whereas areas chosen a lot versus rarely chosen might differ less. In order to make models with more explanatory power to better clarify how the bison uses their available habitat, it would be preferable to work with explanatory variables which gradients overlapped to a lesser extent.

Generally, our results indicate that topography should be considered when choosing or designing rewilding areas. Previous studies looking into habitat use or selection and food selection have not incorporated terrain in their analyses (e.g. 3, 4). It would be interesting to see if habitat preference for drier areas is supported in other reintroduction sites, this might shed more extensive light on the consequences of using European bison as grazers on riparian systems and wet meadows, in addition to dry meadows and forests. In line with Schneider (40) we think that in order to further nature conservation, as well as the conservation of European bison, assessments of how different areas are suited for the bison, and how bison affect the area, plant composition and diversity are needed.

## 5. Conclusion

Overall the present study found similar daily and seasonal behavioural patterns and habitat selection of a semi-free-ranging bison herd as seen in studies on free-living herds, despite the small living space and confinement and a high degree of human modification. These results indicate that rewilding with European bison in anthropogenic environments can support what we, so far, believe is natural behaviour in the wild. This is highly relevant as bison increasingly are being reintroduced to confined enclosures in anthropogenic landscape in order to 1) help conserve the species and 2) restore ecological functions of the ecosystem, which otherwise might be compromised if bison did not express natural behaviour.

However, the unexpectedly high use of the release site compared to the rest of the enclosure prompt us to raise awareness of the possible long-term ethological effects of the release site and the management protocols accomplished here, as this might cause the ecological impact by bison to be reduced in terms of feeding and non-feeding activities. Further, this might reduce the potential of the bison to affect plant diversity, structure and composition, and thereby species communities dependent on these, and thereby compromise or oppose the conservation goals addressed in a trophic rewilding context. We, therefore, encourage managers to match the management in the release area with the overall aims of reintroducing bison in the particular reintroduction area.

European bison is classified as vulnerable by IUCN and therefore still more populations are needed. Combining large-scale rewilding and bison reintroduction might well be a win-win situation, as large-scale rewilding can support bison population and help ensure genetic diversity, and bison can contribute functionally to rewilding projects especially when framed within a trophic-rewilding context. However, still more research is needed to determine optimal habitat of the European bison and to properly assess to what degree European bison thrive in riparian meadows and can replace conventional management protocols such as mowing and cutting regimes.

## Acknowledgments

We are grateful to Randers Regnskov and AVJF for providing us with the opportunity to conduct this study. This work was supported by Aarhus University and the Aage V. Jensen Foundations (PBMP). We also consider this study a contribution to JCS´s Carlsberg Foundation Semper Arden project MegaPast2Future (grant CF16-0005), to the Danish National Research Foundation Niels Bohr professorship project Aarhus University Research on the Anthropocene (AURA), and to JCS´ VILLUM Investigator project (VILLUM FONDEN, grant 16549).

## Supporting information captions

**Table S1.** Multicollinearity of predictor variables considered for use in spatial linear and logistic models. A correlation (r value) of 0.60 was considered a threshold for including variables in the spatial model analyses.

**Figure S1.** Daily average temperature of spring, summer, and fall. Different letters indicate statistical significant differences in daily average temperature among season.

**Figure S2.** Evaluation of spatial autocorrelation based on Moran´s I of first 20 distance classes of model residuals of simultaneous autoregressive lagged (SAR lag) models, simultaneous autoregressive error (SAR err) models, Ordinary Least Square (OLS) regression and Generalized Linear Models (GLM).

